# Simple methods to obtain a food listing and portion size distribution estimates for use in semi-quantitative dietary assessment methods

**DOI:** 10.1101/637025

**Authors:** Christine Hotz, Lubowa Abdelrahman

## Abstract

Semi-quantitative dietary assessment methods are frequently used in low income countries, and the use of photographic series for portion size estimation is gaining popularity. However, when adequate data on commonly consumed foods and portion sizes are not available to design these tools, alternative data sources are needed. This study aimed to develop and test methods to: (i) identify foods likely to be consumed in a study population in rural Uganda, and; (ii) to derive distributions of portion sizes for common foods and dishes. A process was designed to derive detailed food and recipe lists using guided group interviews with women from the survey population, including a ranking for the likelihood of foods being consumed. A rapid recall method to estimate portion sizes using direct weight by a representative sample of the survey population was designed and implemented. Results were compared to data from a 24 hour dietary recall. Of the 82 food items reported in the 24 hour recall survey, 87% were among those ranked with a high or medium likelihood of being consumed and accounted for 95% of kilocalories. Of the most frequently reported foods in the 24 hour recall, portion sizes for many (15/25), but not all foods did not differ significantly (p<0.05) from those in the portion size estimation method. The percent of portion sizes reported in the 24 hour recall between the 5th and 95th percentiles determined by the portion size distribution estimation method ranged from a low of 18% up to 100%. In conclusion, a simple food listing and ranking method effectively identified foods most likely to occur in a dietary survey. A simple method to obtain reliable portion size distributions was effective for many foods, while the approach for others should be modified. These methods are an improvement on those in current use.

## Introduction

Dietary assessment surveys are necessary to adequately inform, design and evaluate nutrition intervention programs in low-income countries. While the 24-hour dietary recall (24HR) is one of the most common method used in these settings [1], it is also more resource intensive and technically challenging than other methods [2,3] and this may be a limiting factor for the use of dietary surveys to inform effective nutrition programs. Depending on the specific objectives of a dietary survey, simpler and less resource intensive methods, including food frequency questionnaires (FFQ) or semi-quantitative (SQ) FFQ, may be adequate [4]. The semi-quantitative estimation of portion sizes consumed using food photo series or atlases depicting graduated portion sizes for a variety of foods is gaining popularity, and may be used to support the application of SQ-FFQs [5] or as a way of simplifying 24HR methods [6,7].

However, to adequately develop such SQ dietary assessment tools, some key information is required beforehand. This includes, but is not limited to: (i) a listing of the foods that are commonly consumed in the study population and hence should be included in the SQ tools; (ii) relevant details about the way they are typically prepared or consumed, and; (iii) the distribution of usual portion sizes to select those to represent in the SQ tools. Ideally, data-driven methods in the form of previously collected dietary intake survey data that is quantitative, valid, and representing the same survey population and sub-population groups of interest, would serve this purpose [5,8]. In low-income countries where data meeting these criteria may often not be available, some form of reliable, empirically-derived preliminary data are needed.

We have found few well-described or well-designed processes in the published literature on how to collect food listing and portion size distribution data when appropriate previous survey data are not available. Food listings have been derived using informal or subjective methods such as consultation with food service professionals, local cook books, or restaurant and cafeteria menus, or conducting interviews with cooks or chefs in households and restaurants, but without any information on the sample size, sampling frame or representativeness [6,9,10]. While these are relatively low-cost methods, it is not clear how complete or representative they are.

In the absence of pre-existing data, portion size ranges have been deduced by various means, such as by adapting from local reference data (e.g., dietary guidelines and nationally established standard serving sizes), consulting experts in the catering industry, qualitative consultation with households [9–11], or deriving a medium portion size from existing survey data but applying fixed ratios to derive small and large portion sizes [9,12]. In some studies to develop and validate food photo atlases, either very scant or no information was provided on how graduated portion sizes were derived [10,13,14], confirming that this methodological step is often overlooked. This is concerning as, if portion size options presented in SQ-FFQ or SQ-24HR dietary assessment surveys do not represent the usual range consumed in the population studied, large portion size estimation errors can result [15].

In some studies, mean portion sizes were determined in small surveys where householders were asked to demonstrate usual portion sizes for different foods for specific age groups [6,12,16], but a minimal description of the methods used for sampling or data collection was provided. This may be an innovative way to collect portion size data for dietary survey tool design, but well-described methods using a systematic and representative approach are needed.

The aim of this study was to develop, document and field test data collection methods to determine food listings and portion size distributions for application in dietary assessment studies. We chose to conduct this work among women in rural Uganda, where researchers have experience in conducting large-scale 24-HR dietary recall and SQ-FFQ surveys. The main objectives were to develop, document, and field test methods to: (i) create listings of foods likely to be consumed in a study population, and; (ii) to quantitatively derive distributions of portion sizes for commonly consumed foods and composite dishes.

## Materials and methods

This study was conducted as part of a larger study to compare dietary intake outcomes of a SQ 24-HR survey method and a SQ-FFQ method with a standard 24-HR method (to be reported elsewhere). The food listing method used a qualitative approach with categorical ranking of the likelihood of foods being consumed, while the method to derive portion size distributions used a quantitative recall approach. The sampling frame was established for the larger study and was used to draw sub-groups of participants for the data collection methods described here. The study was conducted in Nakisunga sub-county (population >48,000) in Mukono District, Uganda, a site that was purposively selected for its proximity to Kampala, having a largely rural agricultural livelihood with some urban influence, socio-cultural homogeneity, and the cooperation of local authorities.

This study was reviewed and approved by the Higher Degrees, Research Ethics Committee, Makerere University School of Health Sciences, Kampala, Uganda and registered and approved by the Uganda National Council of Science and Technology. Informed written consent was obtained from all participants.

## Study participants and sampling

The study included women 18-49 years of age, who were primary residents of the home visited, self-identified as the primary or most senior female caretaker in the household with responsibility for meal preparation, and were available and consented to participate. Women who self-reported to be currently pregnant or lactating with a child <23 months of age were excluded.

We used a multi-stage sampling procedure whereby four of eight parishes in Nakisunga subcounty were randomly selected. Three enumeration areas (EAs), defined by the 2014 Uganda Population and Housing Census sampling frame, were randomly selected from each of the four parishes. A household census of these EAs identified households with eligible women. After dividing the total sample equally among the four parishes, population proportionate sampling of eligible women was done from the three selected EAs for each of the data collection activities. Based on the sample size calculation for the larger study, the samples for these studies were drawn from a pool of n=336 women.

## Data collection

### Socioeconomic data

A brief socioeconomic questionnaire was administered to all participants of the larger study. We used the Progress out of Poverty Index^®^ (PPI), as validated in Uganda, to compare poverty risk among participants in different data collection activities. The questions, indicators and scoring methods were downloaded from the PPI website (http://www.progressoutofpoverty.org/country/ugaada).

### Food listing

The food listing activity was designed to create a list of all foods and beverages commonly consumed by the survey population, including nutritionally-relevant details such as state (e.g., raw/cooked; ripe/unripe), processing method (e.g., dried, milled/extracted, fermented), or cooking method, and to capture main and optional ingredients of commonly consumed recipes.

Two types of semi-structured interviews were used to elicit these details; one with key informants (KI) and the other with groups of survey participants. Prior to the interviews, project staff created a spread sheet with food group categories and an initial list of foods likely to appear.

Two KI interviews were held, one with district level and one with sub-county level government staff. Each interview included a government agriculture officer and a health officer, selected for their knowledge of the availability of local foods, diets, and seasonal availability. A field coordinator and field staff member conducted the interview using the initial food group/food item list as a guide and as a prompt list for foods not mentioned by the KIs, while another field staff member recorded the information. The purpose of the interview, the information of interest, and the ranking categories for likelihood of food items being consumed at the time of the prospective dietary survey (i.e., July 2017) were explained. Detailed recipe data were not obtained in the KI interviews. Data collected from the two interviews were transcribed into a spreadsheet format and combined, and average rankings were calculated, always rounding to the higher likelihood ranking when rankings did not conform. An expanded listing of available food items and common dishes, an average ranking of the of food item availability, and the main processing methods, was derived. This listing was then used to guide the group interviews.

Four guided group interviews (GGI) were held with women randomly selected from the sample list (8-10 women from each of 4 parishes), where each interview covered half of the food groups and lasted approximately 2.5-3 hours. A structured guide was used to elicit food items consumed, including specific details on the food type (e.g., local name(s), color, variety, commercial products), processing and preparation methods (e.g., whole or milled; mashed or chopped and boiled, steamed, fried, etc.), the likelihood of the food being consumed in the household during the survey period (i.e., high, medium, low, not likely at all), and recipes for mixed dishes prepared. For each mixed dish recipe type mentioned, additional details were obtained, including preparation method, whether ingredients were major or minor components, and a likelihood ranking for their inclusion in the recipe (i.e., ‘always’, ‘often’, or ‘rare’; this information is used to correctly identify recipes and for the purpose of collecting standard recipe data, a process not reported on here). Finally, a listing of the most common ingredient combinations was obtained. The data collection tools with completed examples used to record and summarize responses for food items/ingredients and mixed dish recipes, with sample data, are given in S1 and S2 Figs, respectively.

Information from the two interviews on the same food groups was combined in a spreadsheet (one for food items and one for recipe data), and an average ranking score of foods or ingredients was obtained, rounding to the higher frequency category.

### Selection of foods for portion size distribution estimation (PSDE)

Foods consumed as individual items (i.e., not as ingredients in mixed dishes) that were ranked with a high or medium likelihood of being consumed (n=43), plus mixed dishes made with those foods (n=24), were included in the PSDE method, for a total of 67 foods. For the selection of individual food items, this included the 43 high or medium ranked foods identified in the food listing interviews, plus 3 that had oil-fried versions and were distinguished, plus 7 processed baked goods items that were not well addressed in the interviews but added by the researchers as they were considered common in the area. From this total of 53 individual food items, 10 were dropped, as one was not found in the market (i.e., apples), one was better estimated as a count than portion size distribution in grams (i.e., hard candies), and 8 were similar to other items and the portion size was not expected to differ between them (i.e., different meat types, and different varieties of sweet potato, amaranth leaves, yams and some bananas). For the mixed dishes, 30 common ones had been identified, but 6 were dropped as portion sizes were expected to be the same for very similar mixed dishes. For some mixed dishes for which primary ingredients are substitutable, a mixed dish ‘type’ was used to represent the variations (e.g., dishes made with similar types of green leaves or common beans were grouped together).

### PSDE method

Usual portion sizes for different foods were determined with participants using interactive interviews. The 67 selected foods were divided into 4 sets of 16-17 items each, and portion size estimation data were collected on separate days for each set. Four subgroups of 56 women (n=224) invited to participate were asked to recall portion sizes for one of the four sets of foods. We calculated sample sizes for a range of different foods using existing portion size data from a dietary survey conducted in central and eastern Uganda using the equation: [Zα/2. δ / E]^2^, where Zα/2 = 1.96 = 95% confidence, δ = known SD and E = acceptable error in measurement units. The error (E) was set at the equivalent of a coefficient of variation of 15%. This resulted in sample sizes ranging from n=13 to 135, and 80% of the 15 sample sizes calculated were n<60. We rationalized that n=56 data points would be adequate for most foods.

The portion size estimation sessions were organized in a central location of each parish. All foods and dishes were prepared by locally hired assistants in the form typically served. The sets of 16-17 foods were divided into 3 separate data collection stations, with one interviewer and one person weighing and recording the portion sizes, with each woman completing data collection at one station before moving to the next one. For each food item, the interviewer prompted the woman to recall if that food was consumed on the previous day, week, or months. If they could not recall the last time they ate that food, or they never eat that food, no information was collected. If they could recall the last time they ate that food, they were asked to estimate the amount consumed. The respondent was then asked to serve up that amount of food from the real foods provided. These amounts were weighed to the nearest gram on a digital dietary scale, recorded, and the weight of the dish subtracted.

Where portion size data were collected with inedible fractions included (e.g., bones in fish, peel and seeds in watermelon), this was also recorded. Edible fractions for those foods was determined separately by weighing a sample of food items, removing inedible fractions, and then weighing the yield of edible amount on dietary scales. Edible yield factors were then applied to the portion sizes recorded to calculate the weight of the edible portion size.

### 24-HR Survey

We used a multiple pass approach based on Gibson and Ferguson [17] with specific methods that were previously described in detail [18]. Group ‘training’ sessions were held in each EA two days before the 24HR interview to explain the purpose of the study, and the methods involved. They were asked to use their own dishes for serving and eating their food the next day to improve visual memory, and instructed on the use of picture charts to mark foods consumed.

Portion sizes of items consumed were estimated using methods specified for each food type. These included life-sized graduated photographs, weighing scales, graduated measuring cylinders and play dough models, or standard weights for foods that are served as units (e.g., boiled egg, bread slice) [17]. Portion sizes recorded accounted for any leftovers that were served but not consumed. If multiple servings of the same food item were reported to be consumed in a single eating occasion (e.g., morning, afternoon, or evening meals or snacks) these amounts were combined to a single portion. All of these proxy measures were later converted to gram weights of the food represented using a set of conversion factors.

### Data management and analysis

The CSDietary program (HarvestPlus/Serpro, 2009), using the CSPro software platform (Serpro, Santiago, Chile), was used for dietary data entry and data processing. All data were entered in duplicate and discrepancies were identified and rectified and distributions of intakes were reviewed for plausibility by examining high and low intakes. All subsequent data management, processing, and analyses were done using Excel (Microsoft Office 2007 for Windows) and SPSS 16.0 and 18.0 for Windows (SPSS Inc., Chicago, IL, USA). For the food listing, the number of foods by likelihood ranking was determined. For the portion size estimation activity, and portion sizes derived from the 24HR survey, descriptive statistics (mean, SD, CV, and 5th, 50th, and 95th percentiles) were calculated for the portion size distributions in grams.

To determine whether the distribution of portion sizes derived from the PSDE method were adequate to capture those reported in the 24HR survey, we: (i) calculated the number (percent) of portions reported in the 24HR survey whose gram weights fell within the 5th and 95th percentiles of weight derived in the PSDE activity. These percentiles are suggested to represent the smallest and largest portion sizes in food photo series [8], and; (ii) compared the distributions using the Mann-Whitney U-Test/Wilcoxan Rank Sum Test for two independent samples with unequal sample sizes drawn from the same population, where p<0.05 indicates a statistically significant difference. For the socio-demographic data, each individual indicator or score was compared between the PSDE method group and the 24HR survey group, as was the final PPI score.

## Results

### Sample and socio-demographic data

The participation rate in the PSDE was 96% (i.e., 214/224). Socio-demographic data were derived for only a subset of 86% (184/214) of the PSDE participants as these data were collected only for those who also participated in a larger study, including the 24HR survey presented here (Table 1). Of the 184 PSDE participants, 57 participated in both the PSDE and the 24HR survey and the data were retained in both groups for this analysis. Results for these subgroups suggest that they were similar in socio-demographic characteristics (Table 1), except (Table 1).

**Table 1.**
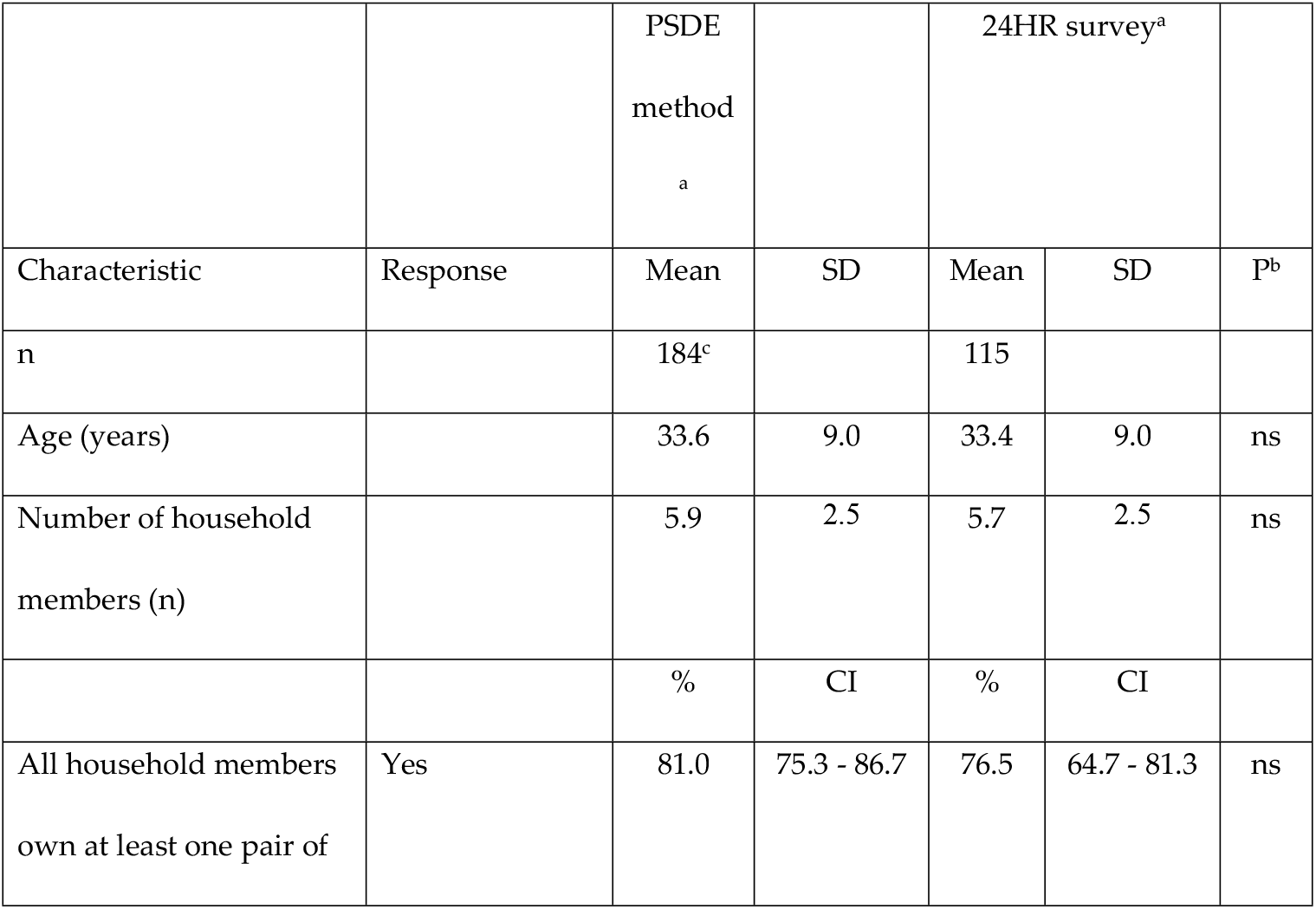
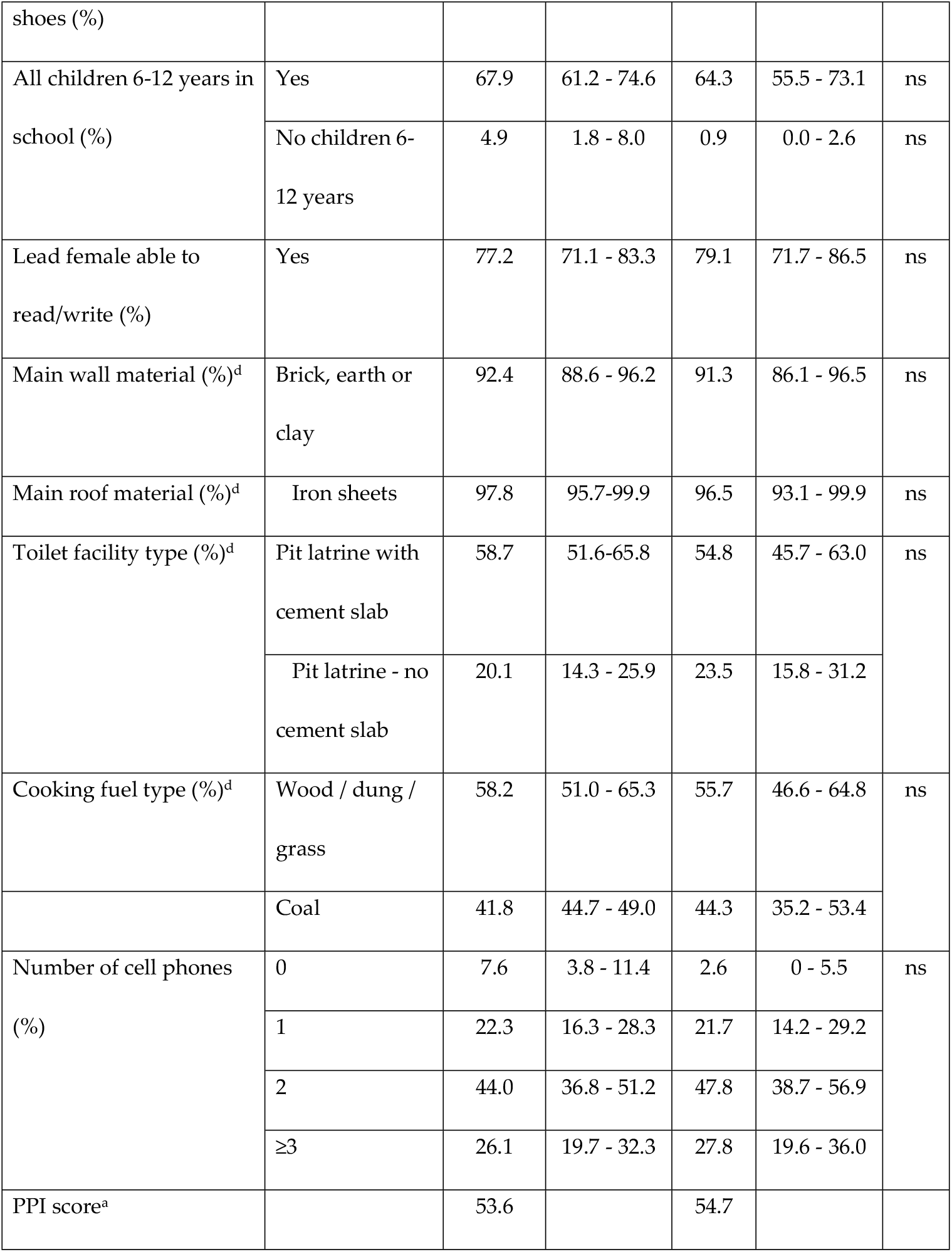
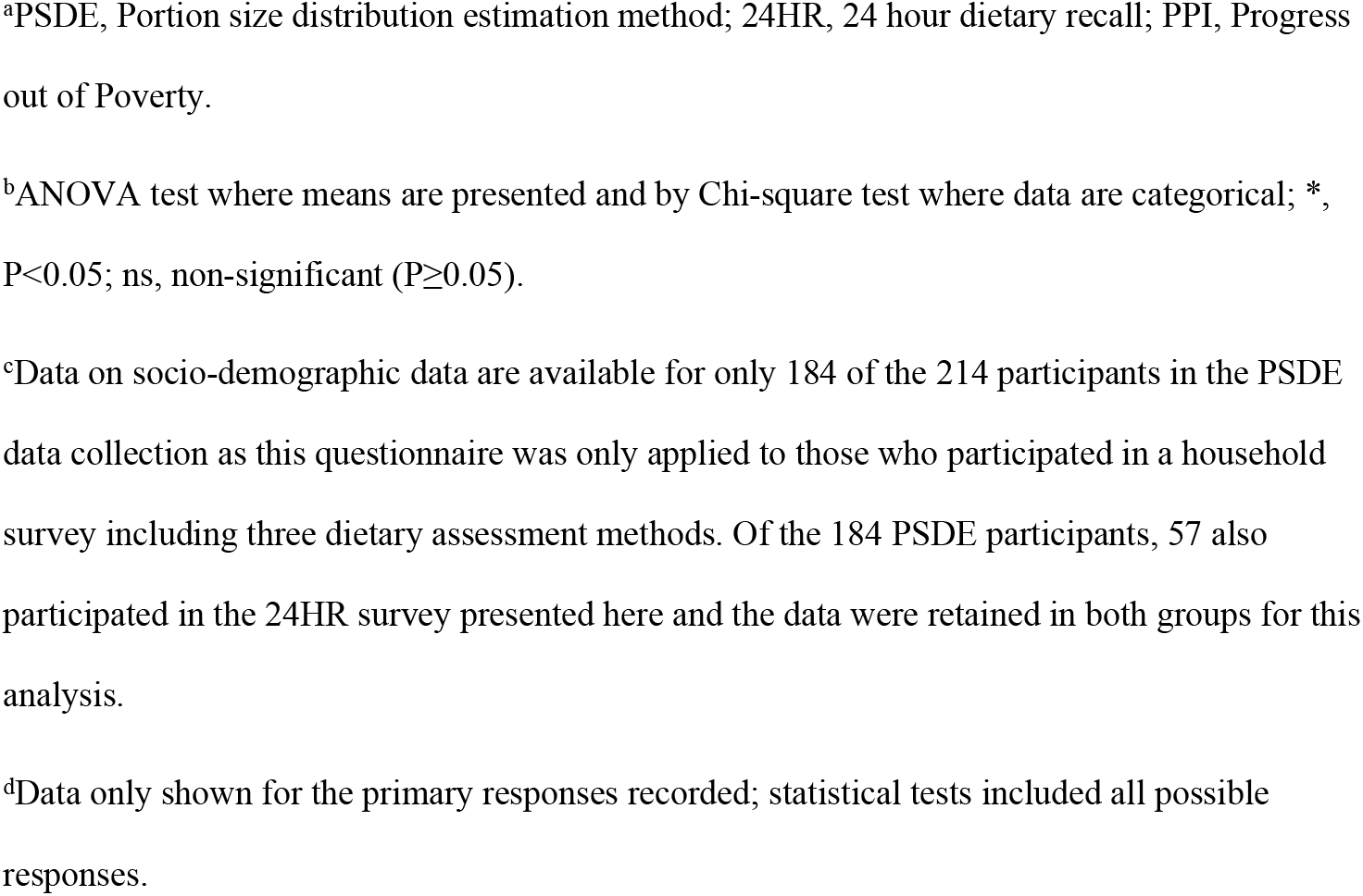
Socio-demographic data for subgroups of participants in the PSDE and 24HR surveys^a^.

|  |  | PSDE<br>method<br><sup>a</sup> |  | 24HR survey<sup>a</sup> |  |  |
| --- | --- | --- | --- | --- | --- | --- |
| Characteristic | Response | Mean | SD | Mean | SD | P<sup>b</sup> |
| n |  | 184<sup>c</sup> |  | 115 |  |  |
| Age (years) |  | 33.6 | 9.0 | 33.4 | 9.0 | ns |
| Number of household<br>members (n) |  | 5.9 | 2.5 | 5.7 | 2.5 | ns |
|  |  | % | CI | % | CI |  |
| All household members<br>own at least one pair of | Yes | 81.0 | 75.3 - 86.7 | 76.5 | 64.7 - 81.3 | ns |
| shoes (%) |  |  |  |  |  |  |
| All children 6-12 years in school (%) | Yes | 67.9 | 61.2 - 74.6 | 64.3 | 55.5 - 73.1 | ns |
|  | No children 6-12 years | 4.9 | 1.8 - 8.0 | 0.9 | 0.0 - 2.6 | ns |
| Lead female able to read/write (%) | Yes | 77.2 | 71.1 - 83.3 | 79.1 | 71.7 - 86.5 | ns |
| Main wall material (%)<sup>d</sup> | Brick, earth or clay | 92.4 | 88.6 - 96.2 | 91.3 | 86.1 - 96.5 | ns |
| Main roof material (%)<sup>d</sup> | Iron sheets | 97.8 | 95.7-99.9 | 96.5 | 93.1 - 99.9 | ns |
| Toilet facility type (%)<sup>d</sup> | Pit latrine with cement slab | 58.7 | 51.6-65.8 | 54.8 | 45.7 - 63.0 | ns |
|  | Pit latrine - no cement slab | 20.1 | 14.3 - 25.9 | 23.5 | 15.8 - 31.2 |  |
| Cooking fuel type (%)<sup>d</sup> | Wood / dung / grass | 58.2 | 51.0 - 65.3 | 55.7 | 46.6 - 64.8 | ns |
|  | Coal | 41.8 | 44.7 - 49.0 | 44.3 | 35.2 - 53.4 |  |
| Number of cell phones (%) | 0 | 7.6 | 3.8 - 11.4 | 2.6 | 0 - 5.5 | ns |
|  | 1 | 22.3 | 16.3 - 28.3 | 21.7 | 14.2 - 29.2 |  |
|  | 2 | 44.0 | 36.8 - 51.2 | 47.8 | 38.7 - 56.9 |  |
|  | ≥3 | 26.1 | 19.7 - 32.3 | 27.8 | 19.6 - 36.0 |  |
| PPI score<sup>a</sup> |  | 53.6 |  | 54.7 |  |  |

<sup>a</sup>PSDE, Portion size distribution estimation method; 24HR, 24 hour dietary recall; PPI, Progress out of Poverty.
<sup>b</sup>ANOVA test where means are presented and by Chi-square test where data are categorical; \*, $P < 0.05$ ; ns, non-significant ( $P \geq 0.05$ ).
<sup>c</sup>Data on socio-demographic data are available for only 184 of the 214 participants in the PSDE data collection as this questionnaire was only applied to those who participated in a household survey including three dietary assessment methods. Of the 184 PSDE participants, 57 also participated in the 24HR survey presented here and the data were retained in both groups for this analysis.
<sup>d</sup>Data only shown for the primary responses recorded; statistical tests included all possible responses.

### Food listing

The food and recipe listing process identified 77 unique foods (i.e., those consumed as individual food items and those used as ingredients in composite dishes) that were ranked in the GGI with a high or medium likelihood of being consumed during the survey, including 3 foods with two distinct preparation methods. Likewise, 48 were ranked with a low likelihood, and 57 as not at all likely or not consumed at all.

Of the 82 distinct foods and ingredients mentioned in the 24HR survey, 71 (87%) were among those ranked with a high or medium likelihood of being consumed and accounted for 95% of estimated kilocalorie intake, while 7% were among those ranked with a low likelihood or not at all likely, accounting for <1% of estimated kilocalories; of the latter, 5 foods were reported by a single individual and 1 food was reported by 2 individuals. The remaining 6% of foods reported in the 24HR did not appear in the food listing at any stage, and accounted for 5% of estimated kilocalorie intake; of these, sugarcane was reportedly consumed by 26 individuals, while the 4 other foods were reported by ≤4 individuals.

The KI food listing tended to result in higher rankings of foods than the FGDs. For example, there were 24 foods ranked as not likely or never consumed by the FGDs that were ranked with a high (n=1), medium (n=8) or low (n=15) likelihood of being consumed. There were also 10 foods listed as being consumed by the KIs but not mentioned or ranked during the GGI. The latter were largely comprised of uncommon bean varieties and non-indigenous vegetables.

### PSDE Method

Descriptive data are presented for the distributions of estimated portion sizes for a selection of individual food items and mixed dishes representing those reported with highest frequency (i.e., >10 occurrences; Table 2) and with low frequency (i.e., 4-6 occurrences; Table 3) in the 24HR survey. The SDs for these food items were relatively large, and the coefficient of variation (CV) for these estimates ranged from 0.27 to 0.98, with an average of 0.47. Portion sizes for approximately half of the individual food items and mixed dishes did not follow a normal distribution (p<0.05).

**Table 2.**
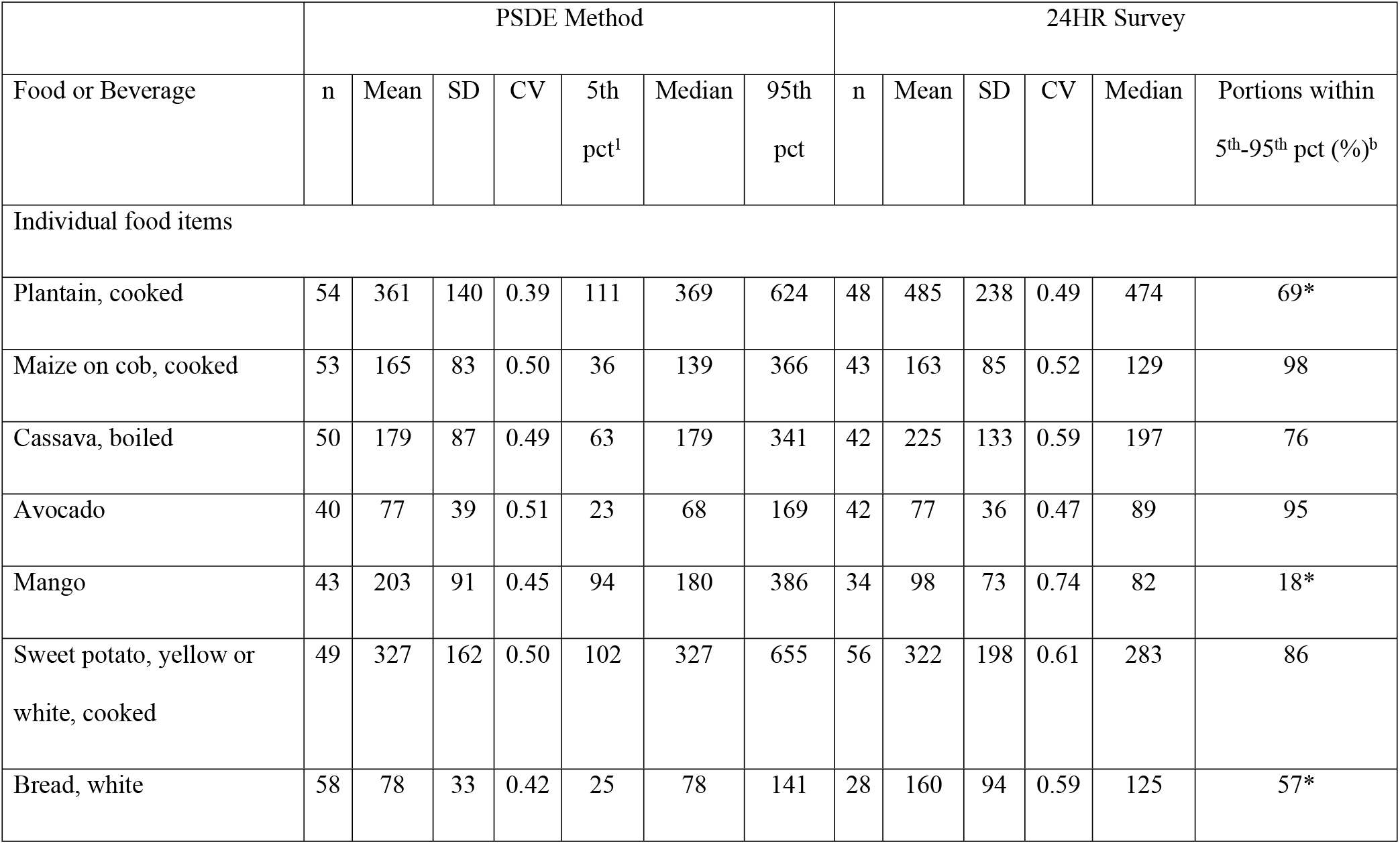
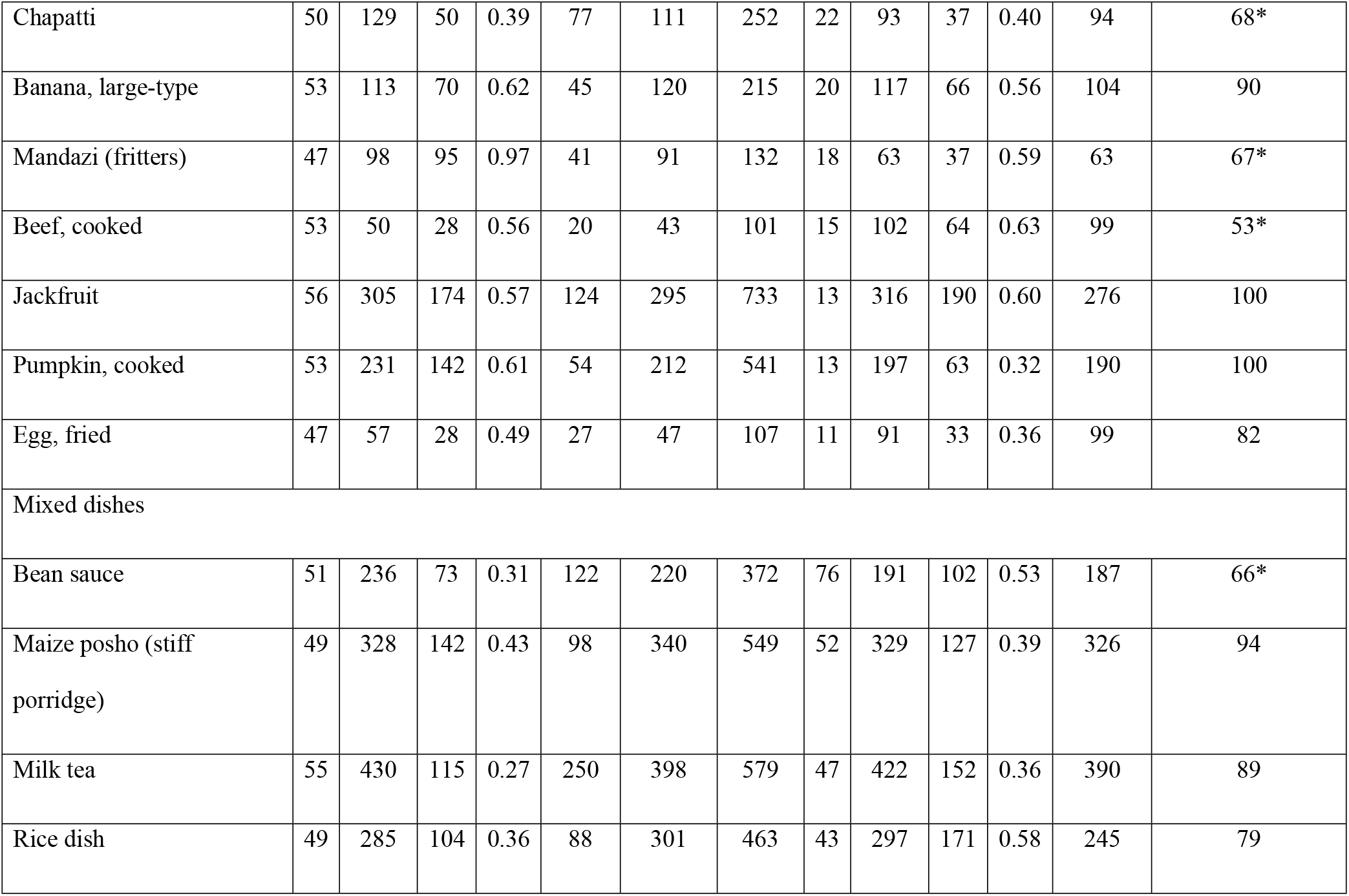
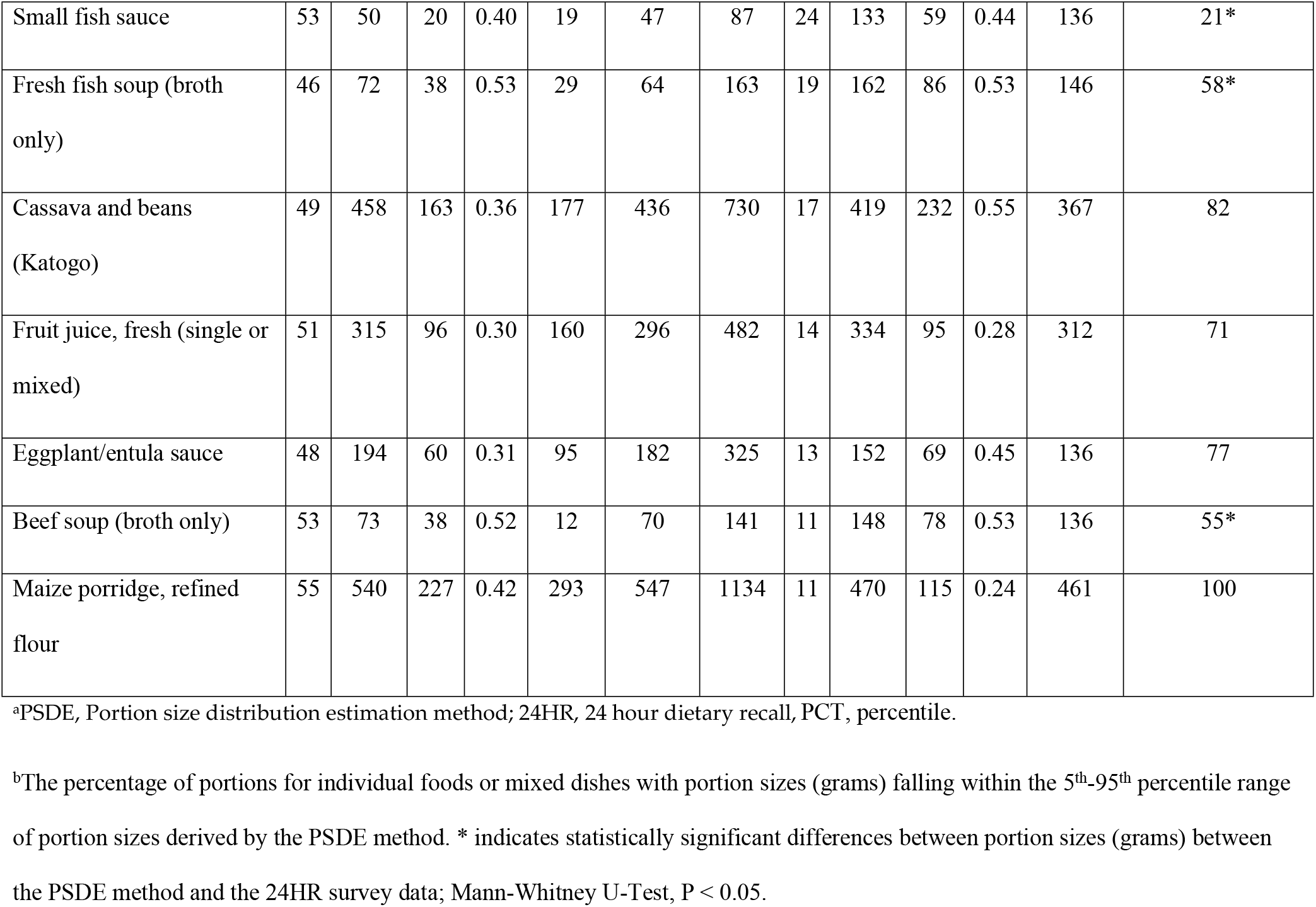
Portion sizes for foods and composite dishes estimated from a portion size recall survey and reported with <u>frequency ≥10% of all food portions</u> in a 24HR survey in the same population.

|  | PSDE Method |  |  |  |  |  |  | 24HR Survey |  |  |  |  |  |
| --- | --- | --- | --- | --- | --- | --- | --- | --- | --- | --- | --- | --- | --- |
| Food or Beverage | n | Mean | SD | CV | 5th<br>pct<sup>1</sup> | Median | 95th<br>pct | n | Mean | SD | CV | Median | Portions within<br>5<sup>th</sup>-95<sup>th</sup> pct (%)<sup>b</sup> |
| Individual food items |  |  |  |  |  |  |  |  |  |  |  |  |  |
| Plantain, cooked | 54 | 361 | 140 | 0.39 | 111 | 369 | 624 | 48 | 485 | 238 | 0.49 | 474 | 69* |
| Maize on cob, cooked | 53 | 165 | 83 | 0.50 | 36 | 139 | 366 | 43 | 163 | 85 | 0.52 | 129 | 98 |
| Cassava, boiled | 50 | 179 | 87 | 0.49 | 63 | 179 | 341 | 42 | 225 | 133 | 0.59 | 197 | 76 |
| Avocado | 40 | 77 | 39 | 0.51 | 23 | 68 | 169 | 42 | 77 | 36 | 0.47 | 89 | 95 |
| Mango | 43 | 203 | 91 | 0.45 | 94 | 180 | 386 | 34 | 98 | 73 | 0.74 | 82 | 18* |
| Sweet potato, yellow or<br>white, cooked | 49 | 327 | 162 | 0.50 | 102 | 327 | 655 | 56 | 322 | 198 | 0.61 | 283 | 86 |
| Bread, white | 58 | 78 | 33 | 0.42 | 25 | 78 | 141 | 28 | 160 | 94 | 0.59 | 125 | 57* |
| Chapatti | 50 | 129 | 50 | 0.39 | 77 | 111 | 252 | 22 | 93 | 37 | 0.40 | 94 | 68* |
| Banana, large-type | 53 | 113 | 70 | 0.62 | 45 | 120 | 215 | 20 | 117 | 66 | 0.56 | 104 | 90 |
| Mandazi (fritters) | 47 | 98 | 95 | 0.97 | 41 | 91 | 132 | 18 | 63 | 37 | 0.59 | 63 | 67* |
| Beef, cooked | 53 | 50 | 28 | 0.56 | 20 | 43 | 101 | 15 | 102 | 64 | 0.63 | 99 | 53* |
| Jackfruit | 56 | 305 | 174 | 0.57 | 124 | 295 | 733 | 13 | 316 | 190 | 0.60 | 276 | 100 |
| Pumpkin, cooked | 53 | 231 | 142 | 0.61 | 54 | 212 | 541 | 13 | 197 | 63 | 0.32 | 190 | 100 |
| Egg, fried | 47 | 57 | 28 | 0.49 | 27 | 47 | 107 | 11 | 91 | 33 | 0.36 | 99 | 82 |
| Mixed dishes |  |  |  |  |  |  |  |  |  |  |  |  |  |
| Bean sauce | 51 | 236 | 73 | 0.31 | 122 | 220 | 372 | 76 | 191 | 102 | 0.53 | 187 | 66* |
| Maize posho (stiff porridge) | 49 | 328 | 142 | 0.43 | 98 | 340 | 549 | 52 | 329 | 127 | 0.39 | 326 | 94 |
| Milk tea | 55 | 430 | 115 | 0.27 | 250 | 398 | 579 | 47 | 422 | 152 | 0.36 | 390 | 89 |
| Rice dish | 49 | 285 | 104 | 0.36 | 88 | 301 | 463 | 43 | 297 | 171 | 0.58 | 245 | 79 |
| Small fish sauce | 53 | 50 | 20 | 0.40 | 19 | 47 | 87 | 24 | 133 | 59 | 0.44 | 136 | 21* |
| Fresh fish soup (broth only) | 46 | 72 | 38 | 0.53 | 29 | 64 | 163 | 19 | 162 | 86 | 0.53 | 146 | 58* |
| Cassava and beans (Katogo) | 49 | 458 | 163 | 0.36 | 177 | 436 | 730 | 17 | 419 | 232 | 0.55 | 367 | 82 |
| Fruit juice, fresh (single or mixed) | 51 | 315 | 96 | 0.30 | 160 | 296 | 482 | 14 | 334 | 95 | 0.28 | 312 | 71 |
| Eggplant/entula sauce | 48 | 194 | 60 | 0.31 | 95 | 182 | 325 | 13 | 152 | 69 | 0.45 | 136 | 77 |
| Beef soup (broth only) | 53 | 73 | 38 | 0.52 | 12 | 70 | 141 | 11 | 148 | 78 | 0.53 | 136 | 55* |
| Maize porridge, refined flour | 55 | 540 | 227 | 0.42 | 293 | 547 | 1134 | 11 | 470 | 115 | 0.24 | 461 | 100 |

<sup>a</sup>PSDE, Portion size distribution estimation method; 24HR, 24 hour dietary recall, PCT, percentile.
<sup>b</sup>The percentage of portions for individual foods or mixed dishes with portion sizes (grams) falling within the 5<sup>th</sup>-95<sup>th</sup> percentile range of portion sizes derived by the PSDE method. \* indicates statistically significant differences between portion sizes (grams) between the PSDE method and the 24HR survey data; Mann-Whitney U-Test, P < 0.05.

**Table 3.** Portion sizes for selected foods and composite dishes estimated from a PSDE survey and reported with low frequency (i.e., 4-6% of all food portions) in a 24HR survey in the same population^a^.

| Data Source | PSDE Method |  |  |  |  |  |  | 24HR Survey |  |  |  |  |  |
| --- | --- | --- | --- | --- | --- | --- | --- | --- | --- | --- | --- | --- | --- |
| Food or Beverage | n | Mean | SD | CV | 5th<br>pct | Median | 95th<br>pct | n | Mean | SD | CV | Median | Portions<br>within 5<sup>th</sup>-<br>95<sup>th</sup> pct<br>(%)<sup>b</sup> |
| Individual food items |  |  |  |  |  |  |  |  |  |  |  |  |  |
| Chips (French fried potatoes) | 40 | 144 | 65 | 0.45 | 56 | 140 | 261 | 5 | 227 | 94 | 0.41 | 260 | 60 |
| Groundnuts, roasted | 50 | 51 | 24 | 0.47 | 20 | 48 | 100 | 5 | 67 | 35 | 0.52 | 67 | 67 |
| Fish (large species), dried,<br>boiled | 49 | 104 | 40 | 0.38 | 43 | 93 | 196 | 5 | 27 | 25 | 0.93 | 21 | 20 |
| Banana, small type | 41 | 148 | 40 | 0.28 | 92 | 142 | 220 | 5 | 160 | 124 | 0.78 | 114 | 20 |
| Samosa | 49 | 64 | 63 | 0.98 | 12 | 61 | 191 | 5 | 95 | 45 | 0.47 | 94 | 100 |
| Mixed dishes |  |  |  |  |  |  |  |  |  |  |  |  |  |
| Plantain & beans (katogo) | 48 | 463 | 154 | 0.33 | 260 | 440 | 764 | 6 | 434 | 136 | 0.31 | 432 | 100 |

<sup>a</sup>PSDE, Portion size distribution estimation method; 24HR, 24 hour dietary recall, PCT, percentile.
<sup>b</sup>The percentage of portions for individual foods or mixed dishes with portion sizes (grams) falling within the 5<sup>th</sup>-95<sup>th</sup> percentile range
of portion sizes derived by the PSDE method.

### 24HR survey

Only 18 individual food items and 11 mixed dishes had ≥10 reported portions consumed, including multiple portions consumed by the same person on the day of recall. Descriptive data (Tables 2 and 3) are not given for 4 individual food items as these were not considered in the portion size estimation activity; two had similar substitute foods included, one was to use a standard unit size as the basis for portion size estimation so was excluded and one food was not picked up in the food listing exercise. For those reported, the SDs were also relatively large and the CVs ranged from 0.24 to 0.93, with an average of 0.49.

### Comparison of portion sizes between the PSE activity and 24HR survey

Of the foods reported with relatively high frequency in the 24HR survey (Table 2), the median portion sizes for many (15/25), but not all foods, were not significantly different from those determined in the PSDE method. For foods with medians that differed significantly, there was no systematic bias in the direction of difference. The percent of portion sizes reported in the 24HR survey that fell between the 5th and 95th percentiles determined by the PSDE method ranged from a low of 18% up to 100%. Of the foods reported with lower frequency in the 24HR survey (Table 3), results were similar. The percentage of portion sizes falling between the 5th and 95th percentiles differed markedly among the 6 food items shown here, ranging from 20 to 100%.

## Discussion

We have described two relatively low cost methods that could aid the development of semi-quantitative dietary assessment methods, as determined in a rural African population. A simple food and recipe listing method found that a ranking system was very effective at identifying foods that were most likely to occur in a dietary survey. A relatively simple and rapid method to obtain reliable distributions of portion sizes from a minimum sample indicated that for many foods, portion size distributions compared well with those obtained from standard 24HR methods, while several others did not.

The food listing method developed and field-tested here provides a useful, categorical method to identify foods that should be included in a food list for dietary surveys using closed lists, such as FFQ and SQ-FFQ methods. The foods identified by the ranking process as having a high or medium likelihood of being consumed covered the vast majority (i.e., 95%) of the total kilocalorie intake in the 24HR survey. This is important as it is a key criteria for developing adequate FFQ/SQ-FFQ methods [5]. If the foods that were ranked here as having a low likelihood or unlikelihood of being consumed were omitted from a FFQ or SQ-FFQ survey derived from it, this would have accounted for a negligible proportion of kilocalories being missed by the survey (i.e., <1%). Although this process may be best suited to general surveys that aim to assess intakes from all foods, it could easily be adapted for use with specific food groups or foods providing specific nutrients.

In addition to use in FFQ and SQ-FFQ surveys, this food listing process could be used to support SQ-24HR methods such as those applying food photo atlases for portion size estimation. It would also be recommended to prepare for standard 24HR surveys as it allows survey designers to create prompt lists for relevant food details that should be probed for in an interview, and to predetermine the most appropriate portion size estimation method for each food likely to occur. This is expected to enhance the training and preparation of enumerators, and possibly the quality of data collected. Very little detail or specific guidance has been provided in the literature where such food listing processes are mentioned [6,9,10,19] or recommended [20].

This food listing method would be improved by including separate likelihood rankings for foods consumed in different forms, including individual foods consumed in raw or cooked forms, foods cooked with or without oil, or as ingredients in mixed dishes so that these can be distinguished for inclusion in the survey. A small number of food items (n=5) occurring in the 24HR survey were not captured by the food listing method. These were primarily low frequency foods, but one food, sugarcane, was reported by a large percentage (i.e., 23%) of individuals.

Baked goods and some commercial beverages were also not adequately probed for during the KIs or GGIs so more careful listing and probing, particularly of processed foods or snacks, is needed, as these can be easily missed under standard food group headings. Finally, the GGI interviews were more relevant to the process as they focused on foods actually consumed in households, rather than focusing more on availability in the community as for the KIs. The latter resulted in more foods being mentioned by KIs and foods being ranked with greater likelihood of consumption than in the GGIs. Nonetheless, the KIs did serve to develop a more complete and locally relevant list of foods for use as a probing guide in the GGIs. The usefulness of combining expert consultation with ethnicity-specific details derived from the target population has been previously recommended [21].

We developed and field tested a novel method to derive portion size distributions for the purpose of developing low cost and simple portion size estimation tools, such as food photo atlases, for use in large-scale dietary surveys. The systematic nature of this method represents an improvement on those reported in the literature for similar use, as previous methods have been largely qualitative in nature, and/or the level of representativeness of the usual range of foods consumed by the target population is questionable [6,9,10,12,14,16].

Portion size estimation tools, including those using photographs, should reflect the range of amounts of foods typically consumed in the study population. A study among children [8] suggested that using age appropriate portion size options greatly reduced error in portion size estimation using photo series depicting portion sizes actually consumed by children (i.e., average of 7% error in weight estimation) compared to using the lower range of portion size photos derived for use with adults (i.e., 46% error) [22]. In an extensive review of FFQ methods [5], it was suggested that in the absence of existing survey data, researchers assigning on the FFQ conduct small surveys to derive portion size data (e.g., 24HR, diet histories). However, in practice this may be impractical due to cost, time, and technical skill required and may not provide reliable distributions where samples for specific foods are less frequent. In the 24HR survey conducted in this study including 111 respondents, the majority of unique foods were reported <10 times, providing a very small sample from which to derive reliable portion size distributions. The method we tested here overcame that problem by quickly obtaining a large sample for each food deemed to be commonly consumed, as identified in the food listing activity.

This study did not aim to validate the PSDE method against a gold standard method, and the comparisons to the 24HR survey must be interpreted cautiously. The PSDE method was limited in that it relied on both short and longer term recall of portion sizes for foods consumed more than a day or week ago, and hence estimates may be distorted by memory. If some responses reflected ‘usual’ portion sizes, the width of distributions may be attenuated as fewer extremes might be reported. This might partially explain why at least some portion sizes reported in the 24HR for most foods were outside the 5th and 95th percentiles of the PSDE distributions.

The 24HR method used different portion size estimation tools, which included photos of small, medium and large items (e.g., vegetables or roots used as ingredients), and dry rice or play dough to estimate volumes, and each of these is then converted to edible portion amounts in grams using previously obtained conversion factors. Thus, some lack of conformity with the PSDE likely occurred due to the difference in methods and additional error that may be introduced by these conversions. In examining results for items for which the distributions were significantly different between methods, there was no apparent bias towards any one portion size estimation tool being consistently associated with low conformity. However, some foods with significantly different distributions had two or more distinct sizes available, such as bread slices from small and large loaves, mandazi (fritters), and small and large mango varieties. In the PSDE method, these were combined into one distribution. However, it’s possible that in the 24HR survey, more individuals were consuming the smaller sizes of those food items and hence the distributions were more skewed to the larger sizes. The lower conformity between methods for standard unit size items (e.g., bread slices, chapatti, mandazi) supports that basing portions on unit size with options for multiples or fractions of those units is a better approach than using continuous portion weights as derived from the PSDE [8]. In the case of beef, the PSDE method accounted for an average amount of bone as part of the beef portions measured, and it’s possible that the 24HR method did not distinguish between meat with or without bone. These are issues that should be considered more deeply in establishing PSE methods and the way foods are included in the PSDE method, as relevant for a particular population.

In addition to supporting the development of FFQs, SQ-FFQ, and simplified 24HRs using food photo atlases for portion size estimation, the PSDE method presented here may also find use in nutrition research and advocacy tools that use linear programming. These methods identify foods that provide, or could provide, sufficient energy or nutrients to meet dietary requirements of a target population and require portion size estimates as input. These include Optifood, primarily used to derive food-based recommendations for optimizing diets of infants and young children [23], and the Cost of the Diet tool, an advocacy tool for estimating the cost of a nutritionally adequate diet [24]. Studies using Optifood typically use 24HR surveys to obtain input data [25,26], while the Cost of the Diet tool does not currently employ a satisfactory method for obtaining usual portion size data on which the models are based; this relatively low-cost method may provide an option to improve this tool.

Validating these methods were beyond the scope of this study. This would require a large-scale quantitative dietary intake survey in the same population with a large enough sample reporting intakes for a wide range of foods. However, we have provided a preliminary, detailed description of methods well beyond what is currently described in the published literature. This includes a limited evaluation comparing to data collected by a quantitative 24HR in the same population, as well as recommendations for improvement. We propose that these methods be further tested and validated when opportunities arise, such as in preparation for a large-scale or national dietary surveys.

## Conclusions

We have identified a gap in available, well-described methods to collect data for deriving food lists and portion size distribution estimates for use in a wide range of dietary assessment methods where existing, suitable dietary intake data are not available. This preliminary evaluation of the methods described and field-tested here, employing qualitative, semi-quantitative and quantitative methods with representative sampling, is encouraging and we recommend efforts to identify the best method of estimating portion size distributions for different food types and to validate these approaches.

## Acknowledgments

The authors are particularly grateful to Carol Nambafu, Doryn Gonahasa, Aisha Sebunya Musaazi, and Vincent Sekajja for their work in supervising and executing the field work in this study, and to Caroline Nankinga and Aisha Namakula at the University of Makerere for their administrative support. We are also grateful to Lynnette Neufeld of the Global Alliance for Improved Nutrition (GAIN), Archileo Kaaya, School of Food Technology, Nutrition and Bio-Engineering, Makerere University and Omar Dary of the United States Agency for International Development (USAID) for their administrative facilitation of the project and technical support in advising on work plans and reviewing project reports.

This study was developed by GAIN and made possible by the generous support of the American people through the support of the U.S. Agency for International Development (USAID) Bureau for Global Health, under the terms of PIO Grant No. GHA-G-00-06-00002. The contents are the responsibility of GAIN and do not necessarily reflect the views of USAID or the United States Government.

## Supporting information captions

**S1 Fig**. Example data collection sheet and guide for food listing in Guided Group Interviews (GGI)

**S2 Fig. Example data collection sheet and guide for detailing recipes of mixed dishes in Guided Group Interviews (GGI)**. We recommend replacing the second column in the second table to record a likelihood ranking for each mixed dish mentioned, rather than obtaining ingredient information.

